# Genomic and proteomic insights into the virulence of a native *Heterorhabditis indica-Photorhabdus akhurstii* complex active against *Spodoptera* spp

**DOI:** 10.64898/2026.10.10.758207

**Authors:** Cecilia Peralta, Nerea Torres-Sanz, Leopoldo Palma

## Abstract

A native entomopathogenic nematode isolate, *Heterorhabditis indica* Che4.1, was recovered from soils in Valencia (Spain), and its symbiotic bacterium, *Photorhabdus akhurstii* Che4.1, was isolated and characterized. Nematode identification was confirmed by ITS and 28S rDNA sequencing, while genome-based analyses supported the taxonomic assignment of the bacterial symbiont. The nematode–bacterium complex exhibited insecticidal activity against *Spodoptera exigua* and *Spodoptera littoralis*. *H. indica* Che4.1 showed an LC₅₀ of 53.7 infective juveniles (IJs)/mL against *S. exigua*, comparable to that of the commercial reference Larvanem^®^ (54.3 IJs/mL). Against *S. littoralis*, Che4.1 exhibited an approximately twofold lower LC₅₀ than Larvanem^®^ (77.3 versus 155.9 IJs/mL), with non-overlapping 95% confidence intervals. The complete genome of *P. akhurstii* Che4.1 consists of a single circular chromosome of 5.66 Mb encoding a diverse repertoire of putative virulence factors, including toxin complex (Tc) components, YenTc-like proteins, Mcf toxins, App-family proteins, Pra/Prb proteins, and chitinases, together with several predicted secondary metabolite biosynthetic gene clusters. LC– MS/MS analyses of bacterial culture supernatants and infected insect cadavers detected multiple predicted virulence-associated proteins, including Tc-related, YenTc-like, App-family, and Pra/Prb proteins. In addition, bacterial cell-free supernatants caused 68.8–81.3% mortality in *S. exigua* neonates under laboratory conditions. Together, these findings reveal a diverse repertoire of potential virulence determinants in *P. akhurstii* Che4.1 and demonstrate the insecticidal potential of its native nematode–bacterium complex. The performance of *H. indica* Che4.1 relative to a commercial reference supports further evaluation of this isolate for biological control of *Spodoptera* spp., particularly under greenhouse and field conditions.

## 1. Introduction

Entomopathogenic nematodes (EPNs) belonging primarily to the genera *Steinernema* and *Heterorhabditis* represent one of the most remarkable examples of mutualistic associations between animals and bacteria used in biological control. These nematodes maintain an obligate symbiosis with gram-negative bacteria of the genera *Xenorhabdus* and *Photorhabdus* (Boemare and Akhurst, 2006), respectively, forming highly specialized pathogenic complexes capable of rapidly killing a wide range of insect hosts in different instars (Del Valle et al., 2017; Del Valle et al., 2016; Tuninetti et al., 2026). During infection, the infective juvenile stage of the nematode actively seeks susceptible insects in the soil environment and penetrates the host through natural openings such as the mouth, anus, or spiracles. Once inside the haemocoel, the nematode releases its bacterial symbionts, which proliferate rapidly and produce a diverse repertoire of virulence factors including toxins, degradative enzymes and secondary metabolites that collectively overcome the insect immune system and lead to host death within 48 h after infestation (Tarasco et al., 2023).

Following host colonization, the bacteria not only contribute to insect mortality but also generate a nutrient-rich environment that supports nematode reproduction and development. At the same time, the production of antimicrobial compounds prevents the growth of competing microorganisms within the insect cadaver. Newly formed infective juveniles subsequently reacquire their symbiotic bacteria and disperse into the environment in search of new hosts, completing a highly efficient biological cycle that underpins the ecological success of these organisms as insect pathogens (Stock and Hazir, 2025).

Because of these characteristics, entomopathogenic nematodes have been widely investigated and commercialized as biological control agents against a variety of agricultural pests, particularly those inhabiting soil environments (Ramakuwela et al., 2025). Their associated bacteria (e.g., *Photorhabdus* spp. or *Xenorhabdus* spp.) are increasingly recognized as rich reservoirs of bioactive molecules, including insecticidal proteins, immunosuppressive factors, degradative enzymes and diverse secondary metabolites with antimicrobial or antiparasitic properties (Cimen et al., 2022; Palma et al., 2024). Recent genomic and metabolomic studies have revealed a remarkable diversity of virulence determinants in *Xenorhabdus* and *Photorhabdus*, including toxin complexes (Tc), Mcf toxins, Pra/Prb binary toxins, pore-forming proteins (Panneerselvam et al., 2022) and numerous biosynthetic gene clusters responsible for the production of biologically active compounds (Cimen et al., 2022; Parihar et al., 2022).

Among lepidopteran pests, species of the genus *Spodoptera* are considered among the most destructive agricultural insects worldwide due to their broad host range, high reproductive capacity and ability to rapidly develop resistance to insecticides (Pogue, 2002). *Spodoptera exigua* and *Spodoptera littoralis* are major pests affecting numerous economically important crops, including vegetables, ornamentals and field crops throughout Europe, Africa and Asia. In recent years, additional concerns have emerged following the expansion of the fall armyworm, *Spodoptera frugiperda*, from its native range in the Americas into Africa, Asia and parts of Europe. The continued spread of this invasive species poses a significant threat to European agriculture and highlights the urgent need for sustainable and effective biological control alternatives capable of targeting multiple *Spodoptera* species.

In parallel, the long-term sustainability of the most widely used microbial insecticide, *Bacillus thuringiensis*, is increasingly challenged by the evolution of resistance in several major agricultural pests (Peralta and Palma, 2017). This situation has stimulated growing interest in alternative or complementary microbial strategies for pest management. In this context, the EPN– bacterium partnership represents a multifactorial pathogenic system fundamentally different from single-toxin approaches such as those based on B. thuringiensis. Unlike Bt-based strategies, which primarily rely on ingestion and are most effective against early larval stages, entomopathogenic nematodes actively seek their hosts and can infect insects through natural openings or directly through the cuticle, allowing them to target pests that are less susceptible to ingestion-based toxins. This infection strategy enables EPNs to kill insects across multiple developmental stages, including late instar larvae and pupae in some species (Del Valle et al., 2017; Tuninetti et al., 2026). Moreover, the combined action of the nematode and its symbiotic bacteria relies on a diverse arsenal of virulence determinants rather than a single toxin, which may reduce the likelihood of resistance evolution. These characteristics, together with their broad host range and compatibility with integrated pest management strategies, position EPNs as promising complementary or alternative microbial agents to conventional Bt-based bioinsecticides.

In this study, we report the isolation and characterization of a native entomopathogenic nematode belonging to the species *Heterorhabditis indica* from soils from a native area in Valencia (Spain) together with its symbiotic bacterium *Photorhabdus akhurstii* strain Che4.1. The bacterial symbiont was isolated, and its complete genome sequenced and analyzed to identify putative virulence factors and biosynthetic gene clusters potentially involved in insect pathogenicity. In addition, the insecticidal activity of both the nematode-bacterium complex and bacterial secreted products was evaluated against two economically important noctuid pests, *Spodoptera exigua* and *Spodoptera littoralis*. These results provide new insights into the pathogenic potential of this EPN-bacterium association and highlight its relevance as a candidate biological control agent.

## 2. Materials and methods

### 2.1. Nematode isolation and characterization

A total of 10 random sub-samples were collected from the Tuéjar riverside at Chelva (Valencia, Spain; 39.7461° N, 1.0061° W). Each sub-sample was taken from a depth of 0-30 cm and combined to form a composite sample of approximately 500 g. The collected soil sample was subsequently sieved to remove debris and moistened (∼10%). The sample was then introduced into 500-cc disposable plastic containers, each containing fourth-instar *Spodoptera littoralis* larvae obtained from our insectary at the Laboratory of Biological Control of Pests (https://cbp.uv.es), University of Valencia. Containers were incubated at room temperature for 7 days. Dead larvae were identified and individually transferred to modified White traps (Kaya and Stock, 1997), which were incubated at room temperature (usually between 7-10 days) until the emergence of the IJs. EPNs were recovered by gentle pipetting and washed three times by centrifugation at 1,500 × g for 5 min using reverse osmosis (RO) filtered water (0.22 µm) and resuspended in 1 mL of water. This nematode suspension was subsequently used to confirm Koch’s postulates using fifth-instar *S. littoralis* larvae and seventh-instar *Galleria mellonella* larvae, also reared in our laboratories.

Three fourth-instar larvae were individually placed in small Petri dishes lined with filter paper, and a non-quantified 0.5 mL of the infective juvenile suspension was applied to ensure contact exposure. Plates were covered and sealed with Parafilm® (perforated to allow gas exchange), incubated at room temperature, and larval mortality was monitored daily. Recovered IJs were washed again as described above and resuspended in dechlorinated filtered water. EPNs were maintained in 4 mL of filtered (0.22 µm) RO water in vented BioLite™ 25 cm² cell-culture treated flasks (Thermo Scientific) to allow adequate aeration. Cultures were stored at 12 °C in darkness in a temperature-controlled wine cooler (Cecotec) and were periodically refreshed or passaged through insect hosts prior to experimental use, or at least every two months to ensure collection viability.

### 2.1. Molecular identification of the nematode

The ITS region was amplified using primers TW81 (5′-GTTTCCGTAGGTGAACCTGC-3′) and AB28 (5′-ATATGCTTAAGTTCAGCGGGT-3′) (Joyce et al., 1994), and the D2–D3 expansion domains of the 28S rRNA gene were amplified using primers D2A (5′-ACAAGTACCGTGAGGGAAAGTTG-3′) and D3B (5′-TCGGAAGGAACCAGCTACTA-3′) (De Ley et al., 1999). PCR reactions were performed in a final volume of 25 μL containing 12.5 μL of NZYTaq II 2× Green Master Mix (NZYTech, Lisbon, Portugal), 0.4 μM of each primer, and 50 ng of genomic DNA, with nuclease-free water added to complete the final volume. Amplifications were carried out under the following cycling conditions: an initial denaturation at 94 °C for 3 min, followed by 35 cycles of denaturation at 94 °C for 30 s, annealing at 56 °C for 30 s, and extension at 72 °C for 1 min, with a final extension at 72 °C for 10 min.

PCR products were separated by electrophoresis on a 1% agarose gel stained with ethidium bromide and visualized under UV illumination. Successful amplicons were purified using the NucleoSpin Gel and PCR Clean-up kit (Macherey-Nagel, Düren, Germany) and sequenced in both directions using the same primers employed for amplification at STAB VIDA (Caparica, Portugal). The obtained sequences were edited and assembled and compared with reference sequences available in GenBank (Sayers et al., 2025) using BLASTn (Altschul et al., 1990).

### 2.2. Symbiotic bacteria isolation and characterization

The symbiotic bacterium was subsequently isolated from *S. littoralis* larval cadavers after surface disinfection with 70% (v/v) ethanol during 1 min. Insect haemolymph was extracted by puncturing the cuticle with a sterile syringe needle, and a droplet was streaked onto Petri dishes containing NBTA agar (37 g nutrient agar, 25 mg bromothymol blue, 4 mL of 0.01 g/mL 2,3,5-triphenyltetrazolium chloride, and 1,000 mL distilled water) supplemented with ampicillin (100 µg/mL). Plates were incubated at 28 °C for 48 h. Colonies exhibiting the characteristic olive-green or bluish coloration typical of *Photorhabdus/Xenorhabdus* were subcultured onto fresh NBTA plates to ensure axenic isolation (Akhurst, 1980). Preliminary molecular characterization was performed by PCR amplification of the 16S rRNA gene, and sequencing as previously described for *Xenorhabdus* strains (Palma et al., 2024).

Total genomic DNA was extracted using the Wizard Genomic DNA Purification Kit (Promega, Madison, WI, USA), following the manufacturer’s protocol for Gram-negative bacteria. DNA integrity was assessed by electrophoresis on 1% agarose gels stained with ethidium bromide (Thermo Fisher Scientific, Waltham, MA, USA), and DNA concentration was quantified using a NanoDrop 2000 spectrophotometer (ThermoFisher Scientific). Purified DNA was used to construct an Illumina sequencing library, which was sequenced at Novogen (UK) using high-throughput Illumina technology. Long-read sequencing was performed using Oxford Nanopore technology at Cucare Diagnostics (Spain) to obtain long DNA fragments, thereby improving genome assembly contiguity and completeness. Raw Illumina and Nanopore reads were quality-checked and trimmed prior to hybrid *de novo* assembly using Unicycler (Wick et al., 2017), integrating short-read accuracy with long-read scaffolding. This approach usually enables the closure of the chromosome and plasmids (if present).

Genome annotation was performed using both the NCBI Prokaryotic Genome Annotation Pipeline (PGAP; 2025 release) and the RAST server (Aziz et al., 2008). Predicted CDS annotated as “Unknown, probable insecticidal toxin CDS” by RAST server, were used to search sequence similarities with the MMseqs2 method at Protein Data Bank (https://www.rcsb.org). In addition, the assembled replicon was also screened using BLAST (Altschul et al., 1990) against a customized, non-redundant insecticidal protein database (2026 update) curated in our laboratory, and candidate hits were subsequently confirmed using BestMatchFinder in the BPPRC database (Panneerselvam et al., 2022).

The circular chromosome map was visualized using the Proksee web server (Grant et al., 2023), allowing the graphical representation of genomic features including coding sequences, GC content and GC skew.

The prediction of putative gene clusters involved in the production of bioactive secondary metabolites was performed with the antiSMASH server (Blin et al., 2019).

Species delimitation and taxonomic assignment were carried out using the Type (Strain) Genome Server (TYGS) (Meier-Kolthoff and Goker, 2019) and PubMLST (Jolley et al., 2018).

### 2.3 Bacterial secretome preparation

The bacterial secretome was obtained from cultures of *P. akhurstii* strain Che4.1 grown in LB broth at 28 °C with shaking at 180 rpm for 48 h. Cultures were centrifuged at 10,000 × g for 10 min at 4 °C to remove bacterial cells. The supernatant was subsequently filtered through a 0.22 µm membrane filter to obtain a sterile cell-free supernatant (CFS). To concentrate extracellular proteins, the CFS was processed using an Amicon® Ultra centrifugal filter unit (Millipore) with a 10 kDa molecular weight cut-off. The retained protein fraction was collected as the concentrated bacterial secretome. Total protein concentration was determined using the Bradford assay (Bradford, 1976), and protein profiles were analyzed by SDS–PAGE to verify the presence of secreted proteins prior to downstream analyses. The concentrated secretome was then stored at −20 °C until further analyses.

### 2.4 LC–MS/MS analysis of the bacterial secretome

Protein identification in the bacterial secretome was performed by liquid chromatography-tandem mass spectrometry (LC–MS/MS). Secretome samples were digested with trypsin and analyzed using a high-resolution LC–MS/MS platform at the Proteomics Facility, Central Service for Experimental Research (SCSIE), University of Valencia, Spain.

Raw mass spectra were processed using MSFragger through the FragPipe platform and searched against a custom protein sequence database derived from the RAST-predicted proteome of *P. akhurstii* Che4.1. Protein identifications were accepted when supported by at least two unique peptides and high-confidence peptide-spectrum matches. Identified proteins were functionally annotated using BLAST and domain prediction tools, and proteins potentially involved in insect pathogenicity (e.g., toxins, chitinases, or degradative enzymes) were highlighted for further analysis.

### 2.5. Proteomic analysis of infected insect cadavers

To identify bacterial proteins produced during infection, freshly killed *S. exigua* larvae infected with *H. indica* Che4.1 were collected and processed for proteomic analysis. Two independent biological replicates were prepared, each consisting of a pool of five dead fourth-instar larvae showing characteristic symptoms of infection after 48 h post infection. Larvae were homogenized on ice in cold phosphate-buffered saline (PBS, pH 7.4) supplemented with 5 mM EDTA and cOmplete™ protease inhibitor cocktail (Roche) at a final concentration of 1×. Homogenates were centrifuged at 16,000 × g for 15 min at 4 °C to remove insoluble material. The resulting supernatants were recovered and filtered through 0.22 µm membrane filters to obtain soluble protein fractions free of cellular debris. Filtered extracts were stored at −80 °C until analysis.

Protein identification was performed by liquid chromatography–tandem mass spectrometry (LC–MS/MS) at the Proteomics Facility, Central Service for Experimental Research (SCSIE), University of Valencia, Spain. Raw spectra were searched against the same custom protein sequence database derived from the RAST annotation of *P. akhurstii* Che4.1. Proteins were accepted when supported by high-confidence peptide-spectrum matches and subsequently annotated according to their predicted biological functions.

### 2.6. Insect Bioassays with EPNs

To evaluate the lethal effects of different nematode doses on insect larvae, one-on-one bioassays were conducted using 24-well polystyrene flat-bottom plates with lids (ThermoFischer) as previously described (Navon and Ascher, 2000). The virulence of *H. indica* Che4.1 was compared with that of Larvanem^®^ (Koppert), a commercial product containing infective juveniles of *Heterorhabditis bacteriophora*, which was included as a commercial reference treatment.

Briefly, a circular piece of filter paper was placed at the bottom of each well, and one fourth-instar insect larva was individually transferred into each well (n = 24 per dose). Nematode suspensions were prepared in water at the designated doses (16, 32, 63, 125, 250, and 500 IJs/mL). A volume of 25 µL of each suspension was applied directly onto each larva, followed by an additional 25 µL of water used to rinse the pipette tip and ensure complete transfer of the infective juveniles. Control larvae received the same total volume of water without nematodes. To prevent larval escape between wells, plates were covered with a layer of absorbent paper prior to closing the lid and incubated at 25 °C under controlled conditions. Larval mortality was recorded every 24 h, and each treatment was evaluated in three independent experiments.

Larval mortality was recorded at 48 h and corrected for control mortality using Abbott’s formula (Abbott, 1925). Dose-response data were analyzed by Probit regression using generalized linear models with a binomial error distribution and Probit link function in R, with nematode concentration expressed on a log₁₀ scale. Median lethal concentrations (LC₅₀) and their 95% confidence intervals were estimated from the fitted models.

### 2.7 Bioassays with bacterial cell-free supernatant

The insecticidal activity of bacterial secreted products was evaluated using cell-free supernatant (CFS). Bioassays were conducted with neonate larvae of *S. exigua* using an artificial diet contamination method.

Three independent bacterial cultures were established from separate colonies and grown in individual Erlenmeyer flasks under identical conditions. Cell-free supernatants were obtained independently from each culture and used as biological replicates (LB1–LB3). Prior to bioassays, the protein concentration of each CFS preparation was determined using the Bradford assay and standardized to 0.65 mg mL⁻¹. A volume of 50 µL of standardized CFS was applied onto the surface of the artificial diet and allowed to dry, corresponding to a final dose of 32.5 µg total protein per diet unit.

Neonate larvae were individually transferred onto the treated diet. Control treatments received sterile LB medium processed in the same manner as the bacterial cultures. Plates were incubated at 28 °C under controlled conditions, and larval mortality was recorded at 24 h intervals for up to 7 days.

Each biological replicate was evaluated using 48 neonate larvae distributed in three independent groups of 16 individuals.

### 2.8 Statistical analysis

To assess the effect of treatment on mortality, a generalized linear model (GLM) with a binomial error distribution and logit link function was fitted using the numbers of dead and surviving individuals as response variables. The significance of treatment effects was evaluated using an analysis of deviance based on likelihood-ratio chi-square tests Estimated marginal means and 95% confidence intervals were obtained from the fitted model. Pairwise comparisons among treatments were performed using Tukey-adjusted contrasts to account for multiple testing.

All statistical analyses were conducted in R version 4.5.3 (R Foundation for Statistical Computing, Vienna, Austria). Statistical significance was considered at p < 0.05.

## 3. Results

### 3.1. Nematode isolation and preliminary characterization

The nematode was able to kill, fifth-instar larvae of *S. littoralis* and seventh-instar larvae of *G. mellonella* in 48 h.

BLAST analysis of both the ITS region and the D2–D3 expansion domains of the 28S rRNA gene consistently identified the isolate as *H. indica*. After sequence curation, the ITS region showed up to 99.7% identity with reference sequences, while the 28S region displayed higher conservation, reaching 100% identity. The obtained sequences have been deposited in GenBank under accession numbers PZ168554 (ITS region) and PZ168575 (D2–D3 region of the 28S rRNA gene).

### 3.2. Isolation and molecular characterization of the symbiotic bacterium

Axenic isolation of the symbiotic bacterium yielded colonies with a characteristic green-olive coloration on NBTA agar plates, consistent with *Photorhabdus* spp. The 16S rRNA gene sequence showed 99.65% pairwise identity with *Photorhabdus luminescens* subsp. *akhurstii* strain IND (Accession No. AY278643). The obtained 16S rRNA gene sequence has been deposited in GenBank under accession number PZ168808.

The genome consists of a single circular chromosome of 5,657,208 bp, with an overall GC content of 42.9%. A graphical representation of the chromosome, including the distribution of predicted virulence genes and biosynthetic gene clusters, is shown in Figure 1. Genome annotation was carried out using the NCBI Prokaryotic Genome Annotation Pipeline (PGAP), predicting a total of 4,911 genes, including 4,638 protein-coding genes and 119 RNA genes. The RNA component comprises 83 tRNA genes, 14 ncRNAs, and multiple copies of ribosomal RNA genes, with 8 copies of 5S rRNA and 7 copies each of 16S and 23S rRNA. In addition, 154 pseudogenes were identified, and four CRISPR arrays were detected in the genome.

**Figure 1.**
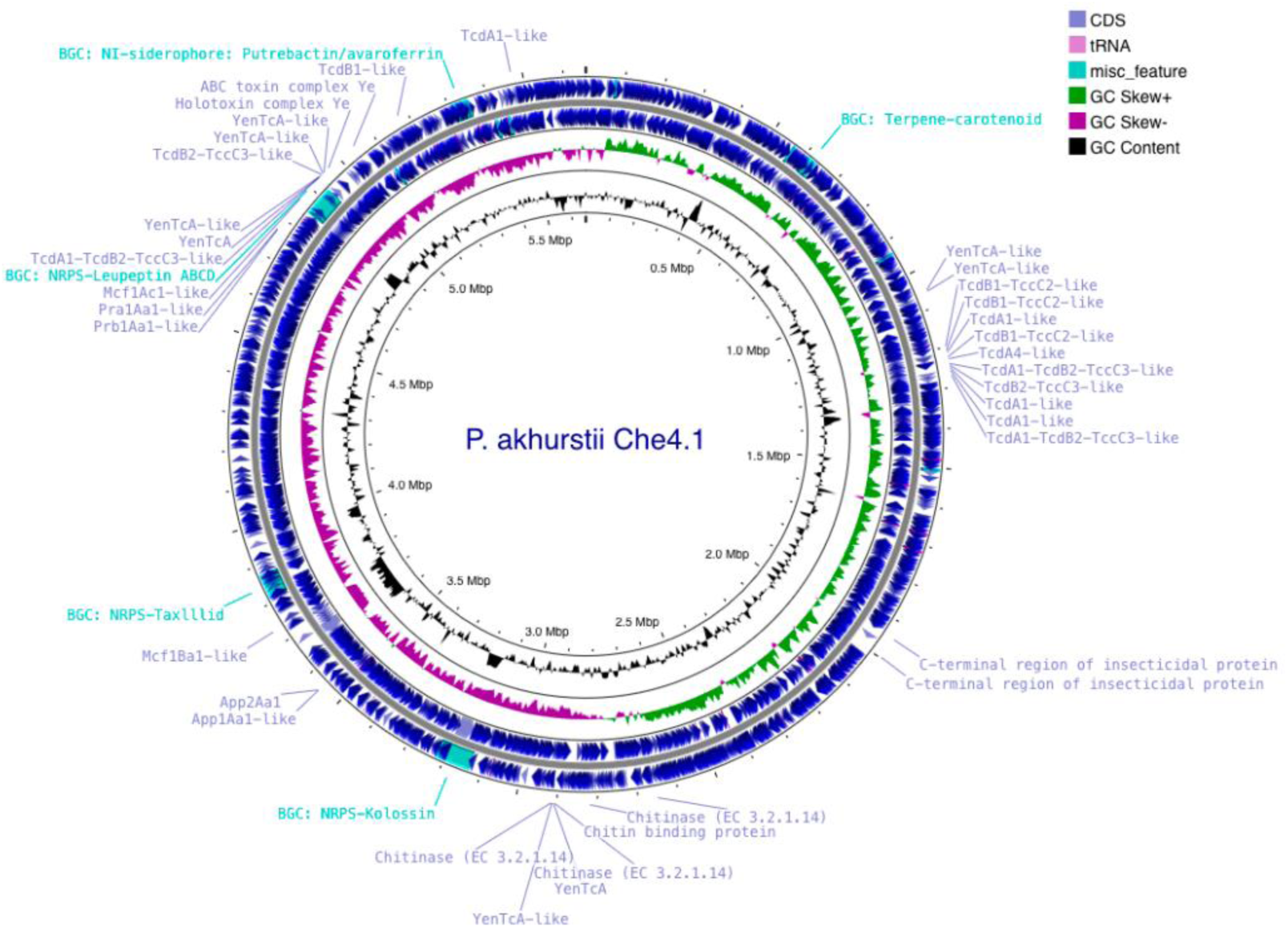
Circular chromosome map of *P. akhurstii strain* Che4.1 generated with Proksee. Coding sequences are represented on the outer rings, with predicted insecticidal protein genes highlighted in blue and biosynthetic gene clusters (BGCs) identified by antiSMASH highlighted in magenta. Inner rings show GC content and GC skew.

Genome-based taxonomic analysis using the Type (Strain) Genome Server (TYGS) confirmed the affiliation of strain Che4.1 within the species *P. akhurstii*. Phylogenomic reconstruction clustered strain Che4.1 with the type strain *P. akhurstii* DSM15138, clearly separating it from other recognized species within the genus *Photorhabdus* (Figure 2). These results support the taxonomic assignment inferred from 16S rRNA gene sequence analysis.

**Figure 2.**
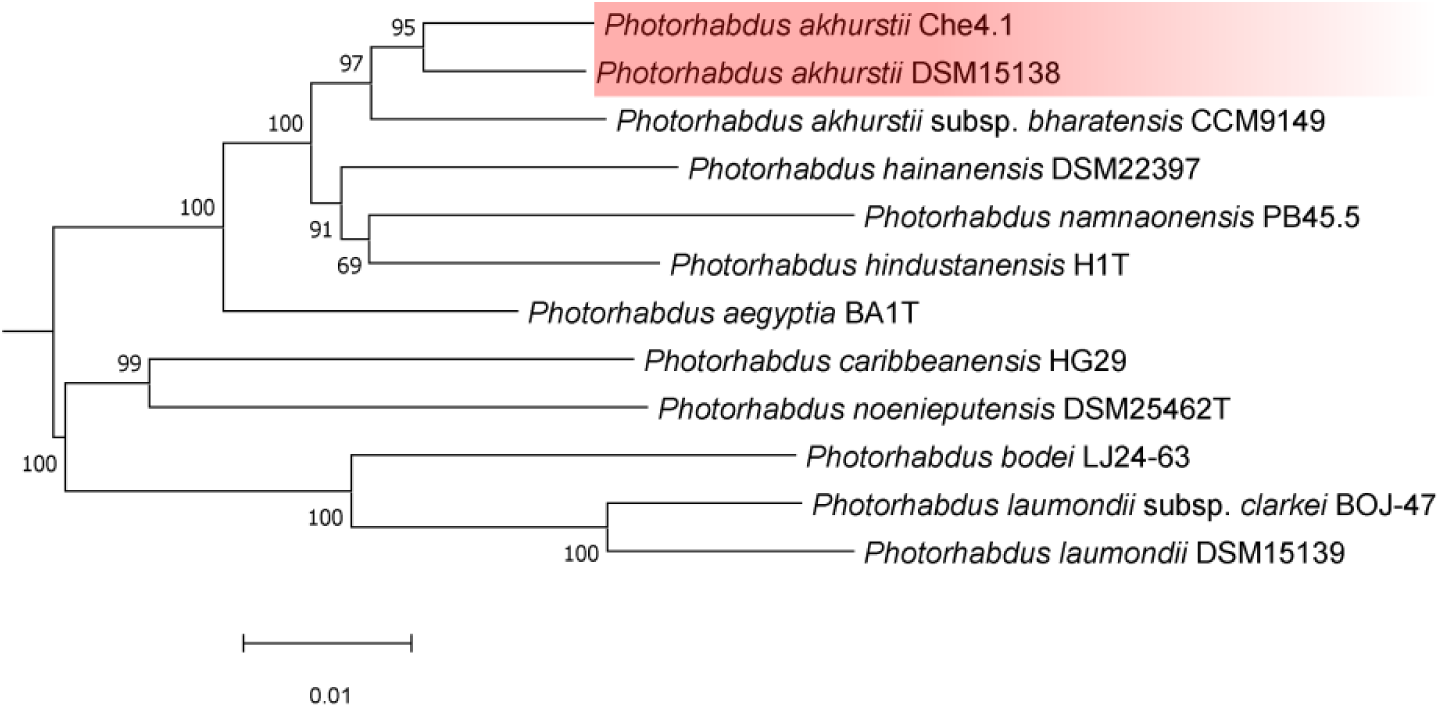
Genome-based phylogenetic placement of strain Che4.1 inferred using the Type (Strain) Genome Server (TYGS). The phylogenomic tree shows the relationship between strain Che4.1 and representative type strains of the genus *Photorhabdus*. Bootstrap support values are indicated at the nodes.

Insecticidal gene annotation revealed genes associated with virulence, mainly multiple putative insecticidal factors and biosynthetic gene clusters. In particular, homologs of well-known *Photorhabdus* insecticidal proteins were identified, including components of the toxin complex (Tc) family (TcdA-, TcdB-, and TccC-like proteins), as well as YenTc-like proteins and Mcf (makes caterpillars floppy)-like toxins. Additionally, genes encoding chitinases and other degradative enzymes were detected. Notably, several of these proteins, including Tc components and chitinases, were also detected in the bacterial secretome by LC–MS/MS analysis (Table 1).

**Table 1.** Putative virulence-associated proteins encoded by the genome of *P. akhurstii* Che4.1 and their detection by LC-MS/MS in cell-free culture supernatants and dead larvae. Predicted proteins include toxin complex (Tc) components, YenTc-like proteins, Mcf toxins, Pra/Prb toxins, chitinases, and other putative insecticidal factors. Protein predictions were based on similarity searches against the BPPRC and Protein Data Bank (PDB) databases.

| Putative virulence factor | ID % | Intervals (bp) | Direction | Homolog Acc. no. | Length (bp) | kDa | Lc-MsMs** |
| --- | --- | --- | --- | --- | --- | --- | --- |
| YenTcA-22 | 25 | 1,051,873-1,054,761 | forward | 6OGD | 2,889 | 109.3 | supernatant |
| YenTcA-14 | 35 | 1,054,809-1,059,206 | forward | 6OGD | 4,398 | 165.4 | supernatant |
| TcdB1-TccC2-27 | 97 | 1,199,934-1,197,187 | reverse | 6L7I | 2,748 | 103.7 | supernatant |
| TcdB1-TccC2-12 | 97 | 1,209,613-1,205,183 | reverse | 6L7I | 4,431 | 165.2 | supernatant |
| TcdA1-4 | 95 | 1,217,219-1,209,666 | reverse | 4O9Y | 7,554 | 282.9 | supernatant+dead larvae |
| TcdB1-TccC2-25 | 83 | 1,225,832-1,223,016 | reverse | 6L7I | 2,817 | 105.1 | supernatant |
| TcdA4-7 | 90 | 1,233,159-1,225,975 | reverse | 6RWA | 7,185 | 204.2 | nd |
| TcdA1-TcdB2-TccC3-20 | 96 | 1,242,805-1,239,893 | reverse | 6H6E | 2,913 | 109.7 | supernatant |
| TcdB2-TccC3-13 | 96 | 1,243,323-1,247,741 | reverse | 4O9X | 4,419 | 165.8 | nd |
| TcdA1-6 | 57 | 1,255,210-1,247,798 | reverse | 8CQ2 | 7,413 | 279.3 | nd |
| TcdA1-8 | 56 | 1,262,217-1,255,684 | reverse | 8CQ2 | 6,534 | 248.2 | nd |
| TcdA1-TcdB2-TccC3-24 | 78 | 1,271,437-1,268,588 | reverse | 6H6E | 2,850 | 106.5 | supernatant |
| C-terminal region of insecticidal protein | - | 1,943,465-1,944,478 | reverse | - | 1,014 | 37.3 | supernatant |
| C-terminal region of insecticidal protein | - | 1,997,226-1,996,183 | reverse | - | 1,044 | 38.3 | supernatant |
| Chitinase 1 | 100 | 2,651,701-2,653,404 | forward | GO:0004568 | 1,704 | 63.9 | supernatant |
| Chitin binding protein | 92 | 2,818,654-2,819,253 | forward | 6T5Z | 600 | 22.4 | supernatant |
| Chitinase 2 | 100 | 2,905,666-2,903,798 | reverse | GO:0004568 | 1,869 | 68.9 | supernatant |
| YenTcA-15 | 69 | 2,909,888-2,905,791 | reverse | 6OGD | 4,098 | 156.3 | supernatant |
| YenTcA-16 | 57 | 2,913,432-2,909,881 | reverse | 6OGD | 3,552 | 133.0 | supernatant |
| Chitinase 3 | 100 | 2,915,164-2,913,530 | reverse | GO:0004568 | 1,635 | 61.4 | supernatant |
| App1Aa1-like (alpha-xenorhabdolysin family) | 84 | 3,555,324-3,556,538 | forward | 6GY6 | 1,215 | 45.7 | dead larvae |
| App2Aa1-like (PaxB) | 98 | 3,556,586-3,557,635 | forward | 6EK4 | 1,050 | 40.3 | supernatant + dead larvae |
| Mcf1Ba1-like (Mcf) | 98 | 3,731,731-3,738,902 | forward | 8P52 | 7,172 | 261.9 | nd |
| Prb1Aa1-like (PirB) | 94 | 4,770,887-4,769,628 | reverse | 7FDP | 1,260 | 46.3 | supernatant + dead larvae |
| Pra1Aa1-like (PirA) | 93 | 4,771,318-4,770,917 | reverse | 7FCN | 402 | 14.8 | supernatant + dead larvae |
| Mcf1Ac1-like (Mcf) | 98 | 4,867,566-4,858,777 | reverse | 8P52 | 8,790 | 324.5 | nd |
| TcdA1-TcdB2-TccC3-18 | 75 | 4,931,345-4,928,214 | reverse | 6H6E | 3,132 | 111.6 | supernatant + dead larvae |
| YenTcA-9 | 32 | 4,936,232-4,931,478 | reverse | 6OGD | 4,755 | 178.6 | supernatant |
| YenTcA-17 | 23 | 4,939,708-4,936,331 | reverse | 6OGD | 3,378 | 122.5 | nd |
| TcdB2-TccC3-19 | 74 | 4,943,760-4,940,629 | reverse | 6SUQ | 3,132 | 111.7 | supernatant |
| YenTcA-10 | 33 | 4,948,588-4,943,894 | reverse | 6OGD | 4,695 | 175.4 | supernatant+dead larvae |
| YenTcA-21 | 58 | 4,951,579-4,948,682 | reverse | 6OGD | 2,898 | 105.5 | supernatant + dead larvae |
| Holotoxin complex Ye-23 | 58 | 4,963,927-4,966,803 | forward | 9C4K | 2,877 | 109.1 | supernatant |
| ABC toxin complex Ye-11 | 60 | 5,044,449-5,048,906 | forward | 4IGL | 4,458 | 166.2 | supernatant+dead larvae |
| TcdB1-26 | 75 | 5,175,491-5,178,301 | forward | 6L7I | 2,811 | 103.9 | supernatant+dead larvae |
| TcdA1-5 | 51 | 5,475,707-5,468,193 | reverse | 4O9Y | 7,515 | 280.7 | supernatant + dead larvae |

Homolog assignments were obtained using BPPRC BestMatchFinder or, when unavailable, inferred from similarity to proteins deposited in the Protein Data Bank (PDB). \*\* A protein was considered detected when its sample-specific spectral count was >0. "Supernatant" indicates detection in at least one culture medium (LB, SY, or TSB); "dead larvae" indicates detection in at least one of the two dead-larva samples; "supernatant + dead larvae" indicates detection in both sample types. nd, not detected by LC-MS/MS in the analyzed samples; -, no homolog identified.

In addition, several biosynthetic gene clusters (BGCs) associated with secondary metabolite production were identified, including terpene, non-ribosomal peptide synthetase (NRPS), and siderophore clusters. Notably, these showed high similarity to known clusters such as carotenoid (pigment production), kolossin (antimicrobial compound), taxlllid (putative bioactive peptide), leupeptin ABCD (protease inhibitor), and putrebactin/avaroferrin (siderophore involved in iron acquisition) (Table 2).

**Table 2.** Predicted biosynthetic gene clusters (BGCs) identified in the genome of *P. akhurstii* strain Che4.1 using antiSMASH analysis. Only clusters showing high similarity confidence to previously characterized BGCs are presented. Predicted clusters include terpene, non-ribosomal peptide synthetase (NRPS), and siderophore-associated pathways potentially involved in antimicrobial activity, iron acquisition, and secondary metabolite production.

| Type | Similarity confidence | Intervals (bp) | Most similar Cluster | Length (bp) |
| --- | --- | --- | --- | --- |
| Terpene | High | 608,475-634,116 | Carotenoid | 25,642 |
| NRPS | High | 3,131,756-3,220,871 | Kolossin | 89,116 |
| NRPS | High | 3,812,037-3,874,752 | Taxllid | 62,716 |
| NRPS | High | 4,850,604-4,926,171 | Leupeptin ABCD | 75,568 |
| NI-siderophore | High | 5,307,824-5,350,296 | Putrebactin/avaroferrin | 42,473 |

\*Only high similarity confidence BGCs are shown.

### 3.3. Nematode bioassays

Both *H. indica* Che4.1 and Larvanem^®^ caused dose-dependent mortality in larvae of *S. exigua* and *S. littoralis* under laboratory conditions. Following correction for control mortality using Abbott’s formula, Probit regression yielded similar LC₅₀ estimates against *S. exigua*: 53.7 IJs/mL (95% CI: 42.0–68.8) for Che4.1 and 54.3 IJs/mL (95% CI: 42.6–69.1) for Larvanem^®^, with extensively overlapping confidence intervals (Table 3). Against *S. littoralis*, Che4.1 exhibited an LC₅₀ of 77.3 IJs/mL (95% CI: 60.8–98.2), compared with 155.9 IJs/mL (95% CI: 118.0–206.1) for Larvanem^®^. Thus, the estimated concentration required to cause 50% mortality in *S. littoralis* was approximately twofold lower for Che4.1 than for the commercial reference, with non-overlapping 95% confidence intervals supporting a difference in potency under the conditions tested.

**Table 3.**
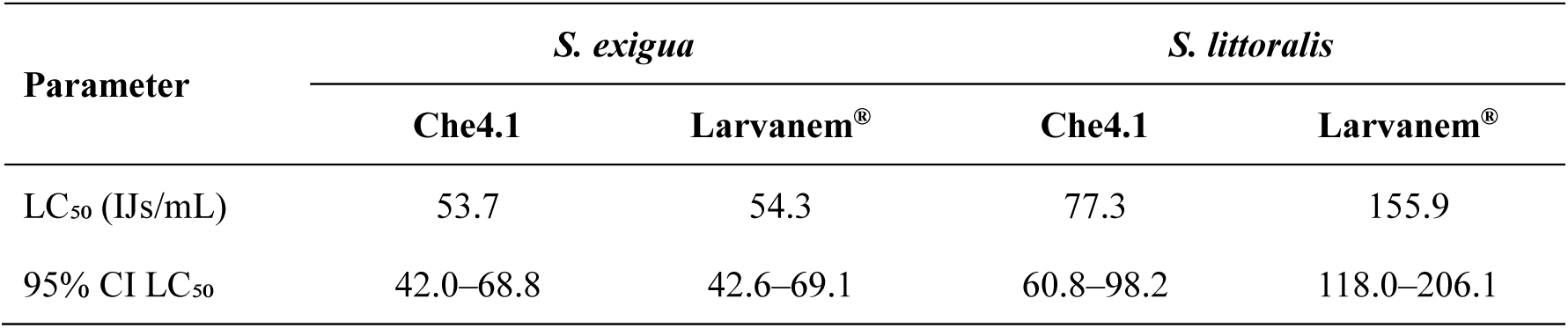
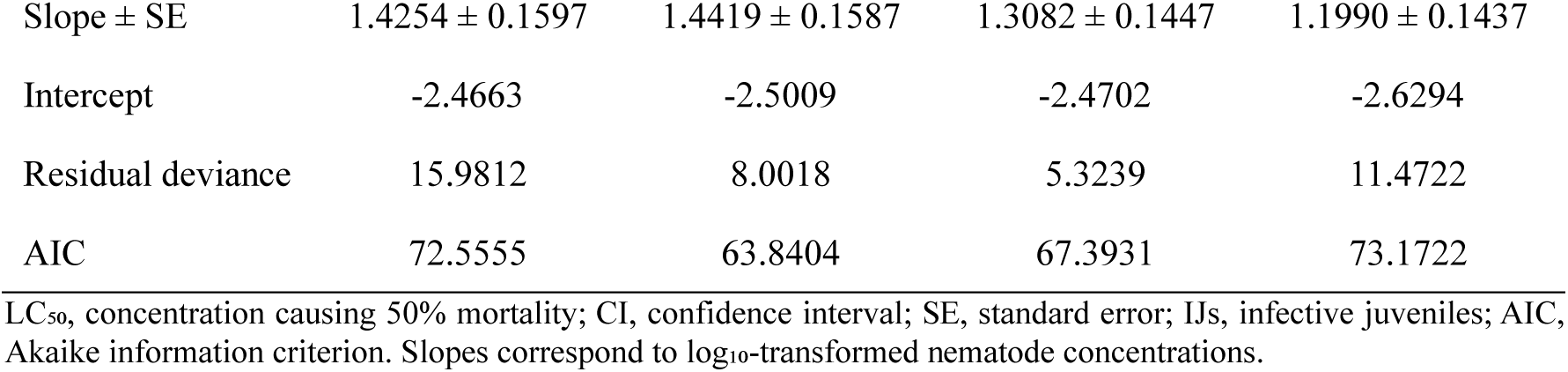
Probit regression parameters and median lethal concentrations (LC₅₀) for *H. indica* Che4.1 and Larvanem^®^ against *S. exigua* and *S. littoralis*, based on mortality corrected for control mortality using Abbott’s formula.

| Parameter | *S. exigua* |  | *S. littoralis* |  |
| --- | --- | --- | --- | --- |
|  | Che4.1 | Larvanem<sup>®</sup> | Che4.1 | Larvanem<sup>®</sup> |
| LC<sub>50</sub> (IJs/mL) | 53.7 | 54.3 | 77.3 | 155.9 |
| 95% CI LC<sub>50</sub> | 42.0–68.8 | 42.6–69.1 | 60.8–98.2 | 118.0–206.1 |
|  | Che4.1 | Larvanem® | Che4.1 | Larvanem® |
| Slope ± SE | 1.4254 ± 0.1597 | 1.4419 ± 0.1587 | 1.3082 ± 0.1447 | 1.1990 ± 0.1437 |
| Intercept | -2.4663 | -2.5009 | -2.4702 | -2.6294 |
| Residual deviance | 15.9812 | 8.0018 | 5.3239 | 11.4722 |
| AIC | 72.5555 | 63.8404 | 67.3931 | 73.1722 |

LC<sub>50</sub>, concentration causing 50% mortality; CI, confidence interval; SE, standard error; IJs, infective juveniles; AIC, Akaike information criterion. Slopes correspond to log<sub>10</sub>-transformed nematode concentrations.

### 3.4. Secretome bioassays

The control treatment showed a mortality of 4.2%, whereas bacterial cell-free supernatants induced mean mortalities of 72.9% (LB1), 68.8% (LB2), and 81.3% (LB3). A generalized linear model (GLM) with binomial error distribution revealed a highly significant effect of treatment on mortality (analysis of deviance: χ² = 83.99, df = 3, p < 0.0001).

Post hoc pairwise comparisons using Tukey-adjusted contrasts indicated that all three supernatant treatments caused significantly higher mortality than the control (LB1 vs Control: p < 0.0001; LB2 vs Control: p < 0.0001; LB3 vs Control: p < 0.0001). In contrast, no significant differences were detected among the three supernatants (LB1 vs LB2: p = 0.970; LB1 vs LB3: p = 0.768; LB2 vs LB3: p = 0.498).

Estimated mortality rates (95% confidence intervals) were 4.2% (1.0–15.2%) for the control, 72.9% (58.8–83.6%) for LB1, 68.8% (54.4–80.2%) for LB2, and 81.3% (67.7–89.9%) for LB3. These results demonstrate that all three cell-free supernatants exhibited significant insecticidal activity against *S. exigua* neonate larvae under the conditions tested, although their efficacy did not differ significantly from one another (Figure 3).

**Figure 3.**
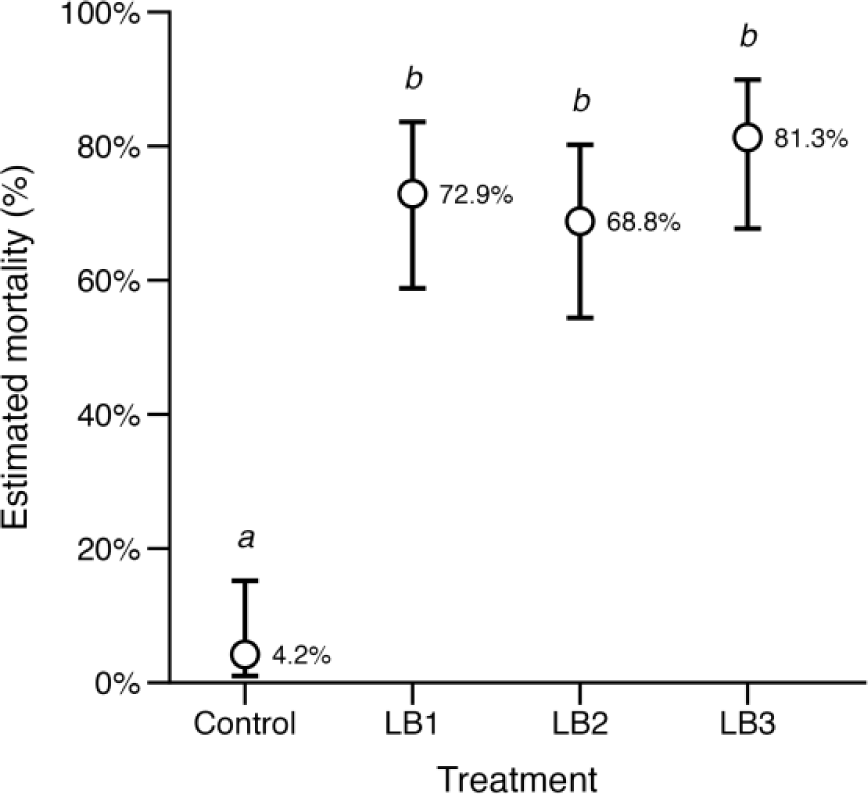
Model-estimated mortality of *S. exigua* neonate larvae following exposure to bacterial cell-free supernatants (LB1–LB3) or the control treatment. Points indicate estimated mean mortality, and error bars represent 95% confidence intervals derived from a generalized linear model with a binomial error distribution. Different letters indicate significant differences among treatments according to Tukey’s multiple-comparison test (P < 0.05).

### 3.5. Proteomic analysis of infected insect cadavers

LC–MS/MS analysis of freshly killed *S. exigua* larvae infected with *H. indica* Che4.1 revealed the presence of multiple bacterial proteins associated with insect pathogenicity. A total of 11 candidate virulence-related proteins were identified (Table 1).

Among the proteins detected in infected cadavers were components of toxin complex (Tc)-related systems, YenTc-like proteins, App-family pesticidal proteins, and Pra/Prb proteins, which have previously been associated with insecticidal activity in entomopathogenic bacteria. An App1Aa1-like protein was detected in infected insect cadavers but not in the analyzed culture supernatants. In contrast, App2Aa1-like, Pra1Aa1-like, and Prb1Aa1-like proteins were detected in both sample types. Additionally, TcdA1-like, YenTcA-like, ABC toxin complex Ye, and TcdB1-like proteins were detected in both bacterial culture supernatants and infected insect cadavers, consistent with their production during infection and suggesting a potential role in insect pathogenicity.

Overall, the proteomic analysis of infected insect cadavers provided evidence for the production of multiple virulence-associated proteins encoded by *P. akhurstii* Che4.1 during infection.

## 4. Discussion

Entomopathogenic nematodes (EPNs) and their associated bacterial symbionts constitute one of the most efficient natural systems for insect pathogenicity and have gained increasing attention as sustainable alternatives to chemical insecticides and conventional microbial control agents. In the present study, we isolated and characterized a native strain of *H. indica* from soils in Valencia (Spain) together with its symbiotic bacterium *P. akhurstii* strain Che4.1, and evaluated their pathogenic potential against *Spodoptera* spp.

Molecular identification based on ITS and 28S rDNA markers consistently assigned the nematode isolate to *H. indica*, a species widely recognized for its broad host range and adaptability to warm environmental conditions. Although *H. indica* has been extensively reported from tropical and subtropical regions, the isolation of a native strain from Mediterranean soils suggests that this species may also be well adapted to local agroclimatic conditions in eastern Spain. Native EPN isolates are often considered advantageous for biological control applications because they may display improved persistence, environmental fitness and compatibility with local ecosystems compared with exotic commercial strains.

The isolated nematode–bacterium complex showed high virulence against *S. exigua* larvae under laboratory conditions, causing rapid mortality within 48 h after infestation. These results reinforce the potential of the native isolate as a candidate biological control agent for lepidopteran pests. In addition, the ability of the isolate to infect different insect species and developmental stages further supports the broad pathogenic potential typically associated with *Heterorhabditis– Photorhabdus* symbiotic complexes.

Comparison with the commercial reference Larvanem^®^ provided a benchmark for assessing the pathogenic potential of *H. indica* Che4.1. The two treatments yielded similar LC₅₀ estimates against *S. exigua*, whereas Che4.1 required approximately half the concentration of infective juveniles to cause 50% mortality in *S. littoralis*. The non-overlapping confidence intervals for the latter comparison support greater potency of Che4.1 against this host under the laboratory conditions tested. This contrasting pattern highlights the importance of evaluating candidate EPN isolates against individual pest species, because their performance relative to a commercial reference may depend on the target host. Although these findings support further evaluation of Che4.1, its efficacy under greenhouse and field conditions remains to be established, including its persistence and performance under different temperature and moisture regimes.

Genome analysis of *P. akhurstii* Che4.1 revealed a large repertoire of putative virulence determinants, particularly toxin complex (Tc)-related proteins, YenTc-like proteins, Mcf toxins, Pra/Prb homologs and multiple chitinases. Similar virulence systems have been previously described in other *Photorhabdus* strains and are known to contribute to host intoxication, immune suppression and tissue degradation during infection. Tc toxins represent one of the most extensively studied insecticidal systems in *Photorhabdus*, functioning as large multi-subunit complexes capable of damaging epithelial tissues and promoting insect mortality. Likewise, Mcf toxins have been associated with apoptosis induction and disruption of insect midgut integrity, whereas Pra/Prb proteins have been linked to oral insecticidal activity in several Gram-negative entomopathogenic bacteria. Particularly noteworthy was the detection of App1Aa1-, App2Aa1-, Pra1Aa1- and Prb1Aa1-like proteins in infected insect cadavers. These proteins were predicted from genome analysis, and their *in vivo* detection provides evidence that they are actively produced during infection, supporting their potential contribution to pathogenicity.

The detection of several Tc-related proteins and chitinases in the bacterial secretome by LC– MS/MS analysis further supports the active production and secretion of virulence-associated factors during bacterial growth. The coexistence of multiple toxin systems together with degradative enzymes suggests that pathogenicity in strain Che4.1 likely relies on a multifactorial mechanism rather than on a single insecticidal determinant. Such combinatorial virulence strategies may provide ecological advantages by facilitating rapid host colonization, nutrient acquisition and evasion of insect immune responses.

The insecticidal activity observed in the cell-free supernatants further supports the functional relevance of the extracellular virulence factors identified in the genome and secretome analyses. All independently produced supernatants caused high mortality of *S. exigua neonates*, whereas the control treatment showed only marginal mortality. The absence of significant differences among biological replicates suggests that the production of insecticidal factors is reproducible under the tested culture conditions. These findings indicate that secreted bacterial products alone are sufficient to induce substantial insect mortality and support the hypothesis that extracellular toxins and degradative enzymes are likely to contribute to the pathogenic process.

In addition to protein toxins, antiSMASH analysis identified several biosynthetic gene clusters potentially involved in the production of bioactive secondary metabolites, including NRPS-associated compounds, siderophores and terpene-related pathways. Some of these clusters showed high similarity to previously characterized pathways involved in antimicrobial activity, protease inhibition or iron acquisition. These metabolites may contribute not only to insect pathogenesis but also to the suppression of competing microorganisms inside the insect cadaver, thereby favoring nematode reproduction and bacterial persistence. The presence of these biosynthetic pathways further highlights the remarkable metabolic versatility of *Photorhabdus* spp. and supports their relevance as reservoirs of biotechnologically valuable compounds.

The increasing emergence of resistance against conventional bioinsecticides, particularly *B. thuringiensis*-based products, has stimulated the search for alternative microbial strategies for pest management. In this context, EPN-associated bacteria provide an attractive source of novel insecticidal molecules with distinct mechanisms of action. Unlike Bt-based approaches, which generally rely on the activity of a limited number of Cry proteins, *Photorhabdus* species possess highly diverse toxin repertoires and secretion systems that may potentially reduce the probability of resistance development in target insects.

Overall, the results obtained in this study highlight the pathogenic potential of the *H. indica–P. akhurstii* Che4.1 complex and provide new genomic and proteomic insights into the molecular basis of virulence in this native isolate. Further studies evaluating field persistence, host range, formulation strategies and the insecticidal activity of purified bacterial metabolites will contribute to determining its applicability as a biological control agent for agricultural pests.

## 5. Conclusions

This study reports the isolation and characterization of a native *H. indica* isolate (Che4.1) from Mediterranean soils in Valencia (Spain) together with its symbiotic bacterium *P. akhurstii* strain Che4.1. The nematode–bacterium complex exhibited high virulence against *S. exigua* larvae under laboratory conditions.

Laboratory bioassays yielded similar LC₅₀ estimates for *H. indica* Che4.1 and the commercial reference Larvanem^®^ against *S. exigua*, while Che4.1 exhibited an approximately twofold lower LC₅₀ against *S. littoralis*. These findings support further evaluation of this locally recovered isolate as a candidate biological control agent for *Spodoptera* pests.

Genome sequencing of *P. akhurstii* Che4.1 revealed a diverse repertoire of putative virulence factors, including toxin complex (Tc)-related proteins, YenTc-like proteins, Mcf toxins, Pra/Prb homologs and multiple chitinases, together with several biosynthetic gene clusters associated with secondary metabolite production. Proteomic analyses of both the bacterial secretome and infected insect cadavers confirmed the production of several virulence-associated proteins during infection.

The coexistence of multiple toxin systems and metabolite biosynthetic pathways suggests that the pathogenicity of the Che4.1 complex relies on a multifactorial mode of action that may enhance insect mortality. These findings reinforce the potential of native EPN-associated bacteria as sources of novel bioactive compounds and support the relevance of the *H. indica–P. akhurstii* Che4.1 association as a promising candidate for sustainable biological control of agricultural pests.

Further studies focused on field performance, formulation development and characterization of secreted insecticidal metabolites will contribute to evaluating its applicability in integrated pest management programs.

## Author Contributions

Conceptualization, L.P.; methodology, C.P.; N.T.-S. and L.P.; validation, C.P. and L.P.; formal analysis, C.P. and L.P.; investigation, C.P.; N.T.-S. and L.P.; resources, L.P.; data curation, L.P.; writing-original draft preparation, C.P. and L.P.; writing-review and editing, L.P.; visualization, L.P.; supervision, L.P.; project administration, L.P.; funding acquisition, L.P. All authors have read and agreed to the published version of the manuscript.

## Data Availability Statement

The complete genome sequence generated in this study has been deposited in GenBank under the accession number CP193895.

## Acknowledgments

Leopoldo Palma gratefully acknowledges the Spanish Ministry of Science, Innovation, and Universities, the Spanish State Research Agency, and the European Union for funding his Ramón y Cajal research contract (grant ref. RYC2023-043507-I). The authors also thank Javier Calvo (Koppert España) for kindly providing Larvanem^®^ for the nematode bioassays.

## References

Abbott, W. S., 1925. A Method of Computing the Effectiveness of an Insecticide. Journal of Economic Entomology. 18, 265–267.

Akhurst, R. J., 1980. Morphological and functional dimorphism in *Xenorhabdus* spp., bacteria symbiotically associated with the insect pathogenic nematodes *Neoaplectana and Heterorhabditis*. J. Gen. Microbiol. 121, 303–309.

Altschul, S. F., et al., 1990. Basic local alignment search tool. J. Mol. Biol. 215, 403–410.

Aziz, R. K., et al., 2008. The RAST Server: rapid annotations using subsystems technology. BMC Genom. 9, 1–15.

Blin, K., et al., 2019. The antiSMASH database version 2: a comprehensive resource on secondary metabolite biosynthetic gene clusters. Nucleic Acids Res. 47, D625–D630.

Boemare, N., Akhurst, R., The Genera Photorhabdus and Xenorhabdus. In: M. Dworkin, et al., Eds.), The Prokaryotes: A Handbook on the Biology of Bacteria Volume 6: Proteobacteria: Gamma Subclass. Springer New York, New York, NY, 2006, pp. 451–494.

Bradford, M. M., 1976. A rapid and sensitive method for the quantitation of microgram quantities of protein utilizing the principle of protein-dye binding. Anal. Biochem. 72, 248–54.

Cimen, H., et al., 2022. Natural products from *Photorhabdus* and *Xenorhabdus*: mechanisms and impacts. Appl Microbiol Biotechnol. 106, 4387–4399.

De Ley, P., et al., 1999. Molecular and morphological characterisation of two reproductively isolated species with mirror-image anatomy (Nematoda: Cephalobidae). Nematology. 1, 591–612.

Del Valle, E. E., et al., 2017. Biological control of *Diloboderus abderus* (Coleoptera: Scarabaeidae) larvae using Steinernema rarum CUL (Nematoda: Steinernematidae) and *Heterorhabditis bacteriophora* SMC (Nematoda: Heterorhabditidae). Crop Protection. 98, 184–190.

Del Valle, E. E., et al., 2016. Biological control of *Alphitobius diaperinus* with *Steinernema rarum* CUL and *Heterorhabditis bacteriophora* SMC and feasibility of application in rice hull. J. Pest. Sci. 89, 161–1701.

Grant, J. R., et al., 2023. Proksee: in-depth characterization and visualization of bacterial genomes. Nucleic acids research. 51, W484–W492.

Jolley, K. A., et al., 2018. Open-access bacterial population genomics: BIGSdb software, the PubMLST.org website and their applications. Wellcome Open Res. 3, 2–20.

Joyce, S. A., et al., 1994. Characterization of *Heterorhabditis* Isolates by PCR Amplification of Segments of mtDNA and rDNA Genes. J Nematol. 26, 260–70.

Kaya, H. K., Stock, S. P. I. L. L. e. M. o. t. i. i. p., Biological techniques series. Academic Press, London, pp 281–324., 1997. Techniques in insect nematology. *In:* Lacey LA (ed) Manual of techniques in insect pathology, Biological techniques series. Academic Press, London, pp 281–324.

Meier-Kolthoff, J. P., Goker, M., 2019. TYGS is an automated high-throughput platform for state-of-the-art genome-based taxonomy. Nat. Commun. 10, 1–10.

Navon, A., Ascher, K., 2000. Bioassays of entomopathogenic microbes and nematodes. Cabi.

Palma, L., et al., 2024. Genome Sequence Analysis of Native *Xenorhabdus* Strains Isolated from Entomopathogenic Nematodes in Argentina. Toxins (Basel). 16.

Panneerselvam, S., et al., 2022. BPPRC database: a web-based tool to access and analyse bacterial pesticidal proteins. Database. 2022, 1–6.

Parihar, R. D., et al., 2022. *Heterorhabditis* and *Photorhabdus* Symbiosis: A Natural Mine of Bioactive Compounds. Front Microbiol. 13, 790339.

Peralta, C., Palma, L., 2017. Is the insect world overcoming the efficacy of *Bacillus thuringiensis*? Toxins. 9, 39.

Pogue, M. G., 2002. A world revision of the genus *Spodoptera* Guenée (Lepidoptera: Noctuidae). American Entomological Society Philadelphia.

Ramakuwela, T., et al., 2025. Entomopathogenic nematodes: Commercial use and future perspectives. Journal of Invertebrate Pathology. 212, 108388.

Sayers, E. W., et al., 2025. GenBank 2025 update. Nucleic acids research. 53, D56–D61.

Stock, S. P., Hazir, S., 2025. The bacterial symbionts of Entomopathogenic nematodes and their role in symbiosis and pathogenesis. Journal of Invertebrate Pathology. 211, 108295.

Tarasco, E., et al., 2023. Entomopathogenic nematodes and their symbiotic bacteria: from genes to field uses. Frontiers in Insect Science. Volume 3–2023.

Tuninetti, F., et al., 2026. Biological control of *Spodoptera cosmioides* pupae by the entomopathogenic nematode *Heterorhabditis bacteriophora* in soybean crops. Phytoparasitica. 54, 34.

Wick, R. R., et al., 2017. Unicycler: Resolving bacterial genome assemblies from short and long sequencing reads. PLoS Comput. Biol. 13, 1–22.

